# Genome report: First whole genome sequence of *Triatoma sanguisuga* (Le Conte, 1855), vector of Chagas disease

**DOI:** 10.1101/2024.10.07.611472

**Authors:** Jennifer K. Peterson, Madolyn L. MacDonald, Vincenzo A. Ellis

## Abstract

*Triatoma sanguisuga* is the most widespread triatomine bug species in the United States (US). The species vectors the human parasite *Trypanosoma cruzi*, which causes Chagas disease. Vector-borne Chagas disease is rarely diagnosed in the US, but *T. sanguisuga* has been implicated in a handful of cases. Despite its public health importance, little is known about the genomics or population genetics of *T. sanguisug*a. Here, we used long-read sequencing to assemble the first whole genome sequence for *T. sanguisuga* using DNA extracted from one adult specimen from Delaware. The final size of the genome was 1.162 Gbp with 77.7x coverage. The assembly consisted of 183 contigs with an N50 size of 94.97 Kb. The Benchmarking Universal Single-Copy Ortholog (BUSCO) complete score was 99.1%, suggesting a very complete assembly. Genome-wide GC level was 33.56%, and DNA methylation was 18.84%. The genome consists of 61.4% repetitive DNA and 17,799 predicted coding genes. The assembled *T. sanguisuga* genome was slightly larger than that of Triatominae species *Triatoma infestans* and *Rhodnius prolixus* (949 Mbp with 90.4% BUSCO score and 706 Mbp with 96.5% BUSCO score, respectively). The *T. sanguisuga* genome is the first North American triatomine species genome to be sequenced, and it is the most complete genome yet for any Triatominae species. The *T. sanguisuga* genome will allow for deeper investigations into epidemiologically relevant aspects of this important vector species, including blood feeding, host seeking, and parasite competence, thus providing potential vector-borne disease management targets and strengthening public health preparedness.

## I. Introduction

Triatomine bugs (‘triatomines’) are hematophagous (i.e., blood-feeding) arthropods that feed on a wide variety of vertebrate host species, including humans. Triatomines are of epidemiological interest due to their harborage of the protozoan parasite *Trypanosoma cruzi*, the causative agent of Chagas disease in humans. If left untreated, infection with *T. cruzi* is lifelong and can lead to serious cardiac and gastrointestinal alterations over time (Rassi Jr *et al*. 2010). There are 162 described species of triatomine bugs (159 extant and 3 fossil species; [Alevi *et al*. 2021; Oliveira Correia *et al*. 2022; Zhao *et al*. 2023; Oliveira-Correia *et al*. 2024]), of which 11 are found in the United States (US; Bern *et al*. 2019).

The most widespread triatomine species in the US is *Triatoma sanguisuga* (Fig. 1). The species has been recorded in 24 states from the southernmost states up to about the 42nd parallel, from the Rocky Mountains to the eastern seaboard (Fig. 2). Spanning over one million square miles, *T. sanguisuga* is found throughout several different ecoregions including most of the great plains, the eastern temperate forests, and the tropical wet forests of southern Florida (United States Environmental Protection Agency 2024).

**Figure 1.**
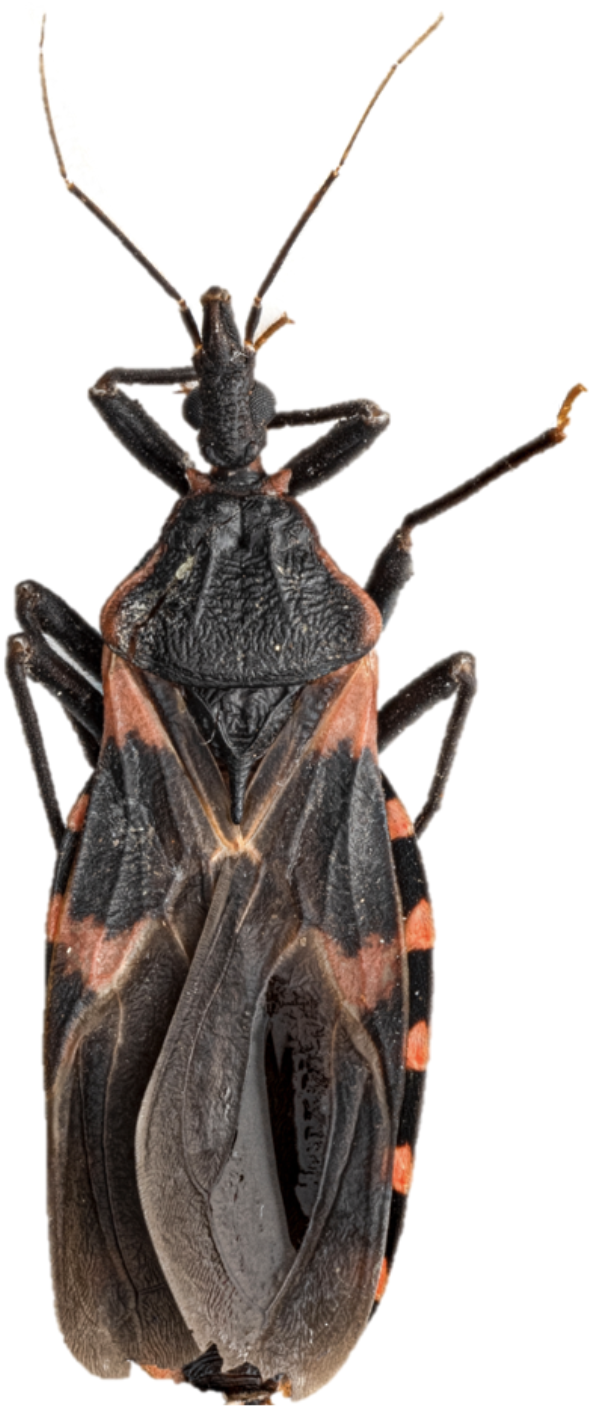
*Triatoma sanguisuga* specimen used in genome sequencing. Image first appeared in figure 2b (Peterson *et al*. 2024). Photo taken by Solomon Hendrix.

**Figure 2.**
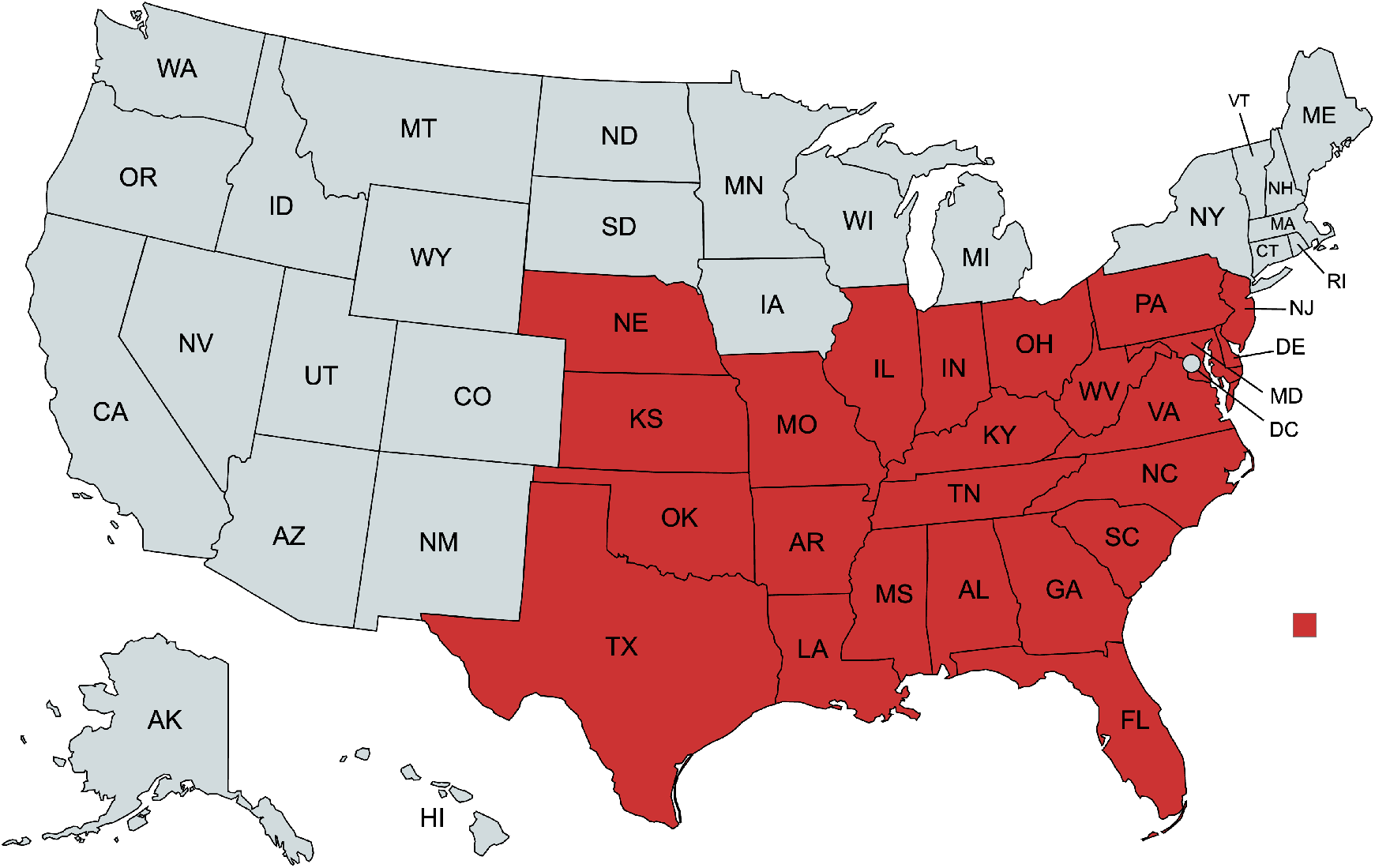
Geographical distribution of *Triatoma sanguisuga*. States with recorded observations of the species are shown in red. Map created with mapchart.net.

*Triatoma sanguisuga* is considered an epidemiologically relevant disease vector species in the US because it can be found in domestic and peridomestic habitats and it has been implicated in autochthonous (i.e., vector-borne) Chagas disease cases (Dorn *et al*. 2007; Lynn *et al*. 2020; Beatty and Klotz 2020). Prior scientific studies of *T. sanguisuga* took place predominantly in three of the 24 states in which the species is found: Louisiana, Texas, and Florida. There, *T. cruzi* infection prevalences upwards of 60% have been found in *T. sanguisuga* (Cesa *et al*. 2011; Moudy *et al*. 2014; Curtis-Robles *et al*. 2018) and blood meal analyses have revealed that the species feeds upon a wide range of taxa comprising reptiles, birds, amphibians and mammals, including humans (Waleckx *et al*. 2014; Dumonteil *et al*. 2020, 2024; Balasubramanian *et al*. 2022).

Despite its ubiquity and epidemiologic importance, little is known about the genetic variation in *T. sanguisuga;* individuals are assigned to the species solely based on morphology and geographic location (De Paiva *et al*. 2022). Five *T. sanguisuga* subspecies have been suggested based on morphological variation, and two of these subspecies were eventually assigned to a new species, *T. indictiva* Neiva 1912. The three remaining subspecies were eventually rejected, but uncertainty still exists (De Paiva *et al*.

2022). Two small studies of *T. sanguisuga* found a high level of genetic diversity between populations. A comparison of cytochrome oxidase II mitochondrial gene sequences from 33 *T. sanguisuga* specimens collected in two barrier islands off the southern coast of Georgia revealed 12 distinct haplotypes with no haplotypes shared between populations from different islands, suggesting limited dispersal and genetic exchange (Roden *et al*. 2011). De La Rua *et al*. (2011) investigated intraspecific genetic variation and population structure in 54 *T. sanguisuga* specimens collected in rural New Orleans, Louisiana and found two groups that were genetically divergent enough to represent different subspecies. The high degree of genetic variation discovered in these two studies hints at a wealth of diversity waiting to be discovered in *T. sanguisuga* considering that its geographic range spans multiple ecological regions with varying seasonality, habitat, and other environmental drivers of genetic diversity.

Here we present the *T. sanguisuga* genome, which is the first genome sequenced for any endemic North American triatomine species. Our whole genome sequencing of *T. sanguisuga* will facilitate comparative analyses between populations to resolve questions of species. In addition, insights into the genetic underpinnings of vector behavior and physiology can lead to new vector control targets. Thus the *T. sanguisuga* genome will contribute to genetic studies of epidemiologically relevant characteristics of *T. sanguisuga* such as blood feeding, host seeking, parasite competence, and domiciliation (Abad-Franch and Monteiro 2005; Fitzpatrick *et al*. 2008; Mesquita *et al*. 2015), and in turn increase our public health preparedness.

## II. Methods & Materials

### Specimen origin

The *T. sanguisuga* specimen used in this study was one of two adult specimens captured within a home in New Castle County, Delaware, as detailed in Peterson *et al*. (2024). The specimens were given to our lab at the University of Delaware by the homeowner. The intestinal contents of both specimens were tested for *T. cruzi* infection via real time PCR as described in Peterson *et al*. (2024). One of the individuals tested positive for *T. cruzi* infection and the other specimen tested negative. The legs and head of the *T. sanguisuga* specimen that tested negative for *T. cruzi* were used for sequencing the genome presented in this study.

### DNA extraction and sequencing

High molecular weight (HMW) genomic DNA extraction and purification from the sample was performed using the MagAttract HMW DNA Kit (Qiagen Inc., Venlo, Netherlands) as per manufacturer’s instructions. High molecular weight DNA was confirmed using a FemtoPulse (Advanced Analytical Technologies Inc., Ankeny, IA). The HMW DNA (10ug aliquots) were converted to SMRTbell templates using the SMRTbell prep kit 3.0 (Pacific Biosciences, Menlo Park, CA) as per manufacturer’s instructions. Briefly, samples were end-repaired and ligated to blunt adaptors. Exonuclease treatment was performed to remove unligated adapters and damaged DNA fragments. Samples were purified using 0.6x AMPureXP beads (Beckman Coulter Inc., Brea, CA). The purified SMRTbell librarieswere eluted in 10 μl of elution buffer. Eluted SMRTbell libraries were size selected on the BluePippin (Sage Science Inc., Beverly, MA) to eliminate library fragments below approximately 10 Kbp. Final library quantification and sizing was carried out on a FemtoPulse (Advanced Analytical Technologies Inc., Ankeny, IA) using 1 μl of library. The amount of primer and polymerase required for the binding reaction was determined using the SMRTbell concentration and library insert size. Primers were annealed and polymerase was bound to the SMRTbell template. Sequencing was performed using the Revio platform (Pacific Biosciences, Menlo Park, CA). The HiFi libraries were run on Revio system 25M SMRT cells using sequencing chemistry 3.0 with 4-hour pre-extension and 30-hour movie time.

### Genome assembly methods

Quality control of the HiFi PacBio reads was performed using NanoPlot v1.43.0 (De Coster and Rademakers 2023). Due to the high quality of the reads, no filtering was needed. The *de novo* assembly of the *T. sanguisuga* genome was performed using Flye v2.9 (Kolmogorov *et al*. 2019), Hifiasm v0.19.5 (Cheng *et al*. 2021), and PacBio’s assembly pipeline in SMRT Link portal v13.1.0 (https://www.pacb.com/smrt-link/). QUAST v5.1.0 (Mikheenko *et al*. 2018) and BUSCO v5.4.7 (Manni *et al*. 2021) were used to assess the completeness of each assembly and compare them to the genome assemblies of two closely related species, *Rhodnius prolixus* (GCA_000181055.3) and *Triatoma infestans* (GCA_011037195.1). BUSCO was run using the ‘hemiptera_odb10’ lineage which consisted of 2,510 single-copy orthologs. The best assembly (the Hifiasm primary assembly) was selected for repeat masking and gene annotation. Full commands for all bioinformatics steps are provided in File S1.

### Decontamination

The Basic Local Alignment Search Tool (BLASTn v2.15.0+; Altschul *et al*. 1990; Camacho *et al*. 2009) was used to query the contigs against the nt_core database v1.1 to check for contamination. Hits with an e-value cutoff of 0.01 and longer than 200 bp were examined. Contigs with only hits to insects and ribosomal RNA were not considered contamination. All other contigs represented possible contamination and were removed from the final assembly and annotation.

### Mitochondria Identification

First, BLAST was used to query the contigs against the existing *T. sanguisuga* mitochondrial sequence (NC_050329.1). Hits were filtered by query coverage to retain contigs with more than 50% of their sequence aligning to the mitochondrial sequence. To confirm the possible mitochondrial contigs and select a representative, MitoHifi v3.2.2 (Uliano-Silva *et al*. 2023) was run via its Singularity container. MitoHifi was also used to annotate and circularize the representative contig and modify it to start at tRNA-Phe. The BLAST alignment between the representative contig and the existing *T. sanguisuga* mitochondrial sequence was used to identify any regions in the representative contig that was not in the reference mitochondrial sequence. Identified significant region(s) were extracted using bedtools and queried against nt and Univec build 10 (https://www.ncbi.nlm.nih.gov/tools/vecscreen/univec/) databases to determine the potential source of the sequence. Any region with no clear source (no significant hits to either database) was removed. The resulting sequence was re-annotated with MitoHifi and visualized using circularMT (Goodman and Carr 2024).

### Methylation

PacBio sequencing enables detection of 5-methylated cytosines (5mCs; Flusberg *et al*. 2010). Hifi reads with the 5mC information (5mC tags in an unaligned bam file) were aligned to the Hifiasm primary assembly using Pbmm2 v1.14.0 (https://github.com/PacificBiosciences/pbmm2), a C++ wrapper for minimap2 v2.26 (Li et al., 2018). PacBio’s pb-CpG-tools v2.3.2 (https://github.com/PacificBiosciences/pb-CpG-tools) was then used to produce per-site methylation probabilities from the alignment. Global CpG methylation was calculated by dividing the number of methylated CpG cytosines by the number of unmethylated cytosines in the reads that aligned to the genome assembly.

### Repeat assembly techniques

Repeats in the Hifiasm primary assembly were identified and modeled using RepeatModeler v2.0.3 (Flynn *et al*. 2020). Repeats from this *de novo* identification were masked using RepeatMasker v4.1.2 (Smit *et al*. 2013). In addition, RepeatMasker was run using Dfam database v3.2 and the parameter ‘-species Triatoma’. Complex repeats were extracted from the RepeatMasker results and given to the Maker annotation pipeline (Cantarel *et al*. 2008; Campbell *et al*. 2014) for hard-masking (see next section).

### Gene finding methods

Annotation was performed using Maker v3.01.04 (Cantarel *et al*. 2008). First, an evidence based round of Maker was run using the transcripts from *R. prolixus* obtained from VectorBase release 68 (Giraldo-Calderón *et al*. 2015) and the ‘Triatominae’ canonical proteins from UniProt release 2024_3 (Apweiler *et al*. 2004; The UniProt Consortium 2023) as the transcript and protein evidence respectively. For the repeat masking parameters of Maker, the ‘rm_gff’ parameter was set to the complex repeats GFF from RepeatMasker, ‘model_org’ was set to ‘simple’, and ‘repeat_protein’ was set to Maker’s provided ‘te_proteins.fasta’ file. The transcripts (with added 1,000 bp flanking regions) from this first round of Maker were then used to train gene models using Augustus v3.5.0 (Stanke *et al*. 2006) via BUSCO v5.4.7 long mode. The starting Augustus species was set to ‘rhodnius’. The new Augustus model was then provided into a second round of Maker to produce the final set of predicted gene annotations. Putative gene function was assigned by following Maker Support Protocols 2 and 3 (Campbell *et al*. 2014) and functional domains and GO terms were assigned via Interproscan v5.53-87.0 (Blum *et al*. 2020) by following steps 4 and 5 of Basic Protocol 5 (Campbell *et al*. 2014). The predicted genes were then sorted into a high confidence set by filtering out genes with no function assigned and an AED of 1 (no transcript or protein evidence from Maker). BUSCO with the ‘hemiptera_odb10’ lineage was used to assess the completeness of the high confidence proteins and transcripts. The quality of the annotated proteins was also examined by identifying their *R. prolixus* orthologs using the OrthoVenn3 web service (Sun *et al*. 2023) with the Orthofinder algorithm (Emms and Kelly 2019). All annotation files are located in file S2.

### Comparative analysis

Whole genome sequences have been assembled and deposited in the National Center for Biotechnology Information (NCBI) for two other species in the subfamily Triatominae, *R. prolixus* (Bioproject ID PRJNA13648) and *T. infestans* (Bioproject ID PRJNA589079). Although now eliminated from the region, *R. prolixus* is believed responsible for the majority of current Chagas disease cases in Central America (Hashimoto 2012; Peterson *et al*. 2019a, 2019b), and it is still one of the main vectors of Chagas disease in Colombia and Venezuela (Fitzpatrick *et al*. 2008; Méndez-Cardona *et al*. 2022). *Triatoma infestans* is found in the southern cone of South America and it is one of the main vectors of Chagas disease cases in that region (Coura, 2014). Note that two additional genomes of *T. infestans* have reportedly been sequenced (Pita *et al*. 2017a, 2017b, 2018; Mora *et al*. 2023), but the data are not publicly available. For quantitative comparison, we used data from the publicly available *T. infestans* genome.

## III. Results and Discussion

### Assembly

A total of 91.08 Gbp of sequence was generated with an N50 of 14,271.0 bp and 92.2% of reads above the Q20 quality cutoff and 74.6% above the Q25 cutoff (Table 1). The Hifiasm assembly was selected for annotation based on a comparison of assembly quality statistics and BUSCO scores for assemblies by the assemblers Flye, Hifiasm, and Smrtlink assemblers, as described above (Table 2). The best assembly was the Hifiasm assembly, which was 1.165 Gbp spread over 282 contigs. Decontamination revealed two contigs not pertaining to *T. sanguisuga*; the first contig aligned (>90% identity) with sequences from mitochondria of three fungal species belonging to the order Hypocreales, which includes several entomopathogenic species. The second contig aligned to a tick-borne protozoan species, *Hepatozoon canis*. Both contigs were removed from the Hifiasm assembly, which decreased the genome size by 105,876 bp (Table 2). Analysis of the mitogenome revealed 98 redundant contigs, 97 of which were removed, resulting in a final assembly of 1.162 Gbp spread over 183 contigs with an N50 of 94,972,618 bp and coverage of 77.7x. The genome-wide GC level was 33.56%. All sequence and assembly data, including sequences of the two contaminated contigs that were removed, are available under NCBI BioProject ID PRJNA1140168 and accession number SRR29988702.

**Table 1.** Quality cutoff data for *T. sanguisuga* sequencing reads.

| Quality cutoff | # reads above cutoff | % of reads above cutoff | Mb of reads above cutoff |
| --- | --- | --- | --- |
| Q10 | 6,532,647 | 100.0% | 91,075.0 Mb |
| Q15 | 6,529,241 | 99.9% | 91,027.9 Mb |
| Q20 | 6,021,510 | 92.2% | 83,537.3 Mb |
| Q25 | 4,872,187 | 74.6% | 66,254.9 Mb |
| Q30 | 3,438,652 | 52.6% | 44,709.0 Mb |

**Table 2.** Quality statistics for assemblies of the *T. sanguisuga* genome by three different *de novo* assemblers. The Hifiasm draft one assembly was selected as the best assembly and thereafter underwent decontamination to produce the draft two assembly, followed by removal of 97 redundant contigs pertaining to the *T. sanguisuga* mitochondria to produce the final assembly.

|  | Flye | Smrtlink | Hifiasm<br>(Draft 1) | Hifiasm<br>Decontaminated<br>(Draft 2) | Final<br>assembly |
| --- | --- | --- | --- | --- | --- |
| **ASSEMBLY** |  |  |  |  |  |
| **Length (bp)** | 2,079,745,197 | 1,180,738,419 | 1,165,190,363 | 1,165,084,487 | 1,162,099,166 |
| **Contigs** | 9,026 | 1,068 | 282 | 280 | 183 |
| **Largest contig** | 7,311,449 | 24,892,402 | 144,379,499 | 144,379,499 | 144,379,499 |
| **Min contig length** | 1,021 | 1,811 | 9,031 | 9,031 | 9,031 |
| **Mean contig length** | 230,417.2 | 1,105,560.3 | 4,131,880.7 | 4,161,016 | 6,350,268.7 |
| **N50** | 388,970 | 6,977,916 | 94,972,618 | 94,972,618 | 94,972,618 |
| **L50** | 1,563 | 52 | 5 | 5 | 5 |
| **BUSCO** |  |  |  |  |  |
| **Complete** | 98.9% | 98.8% | 99.1% | 99.1% | 99.1% |
| **Single** | 36.4% | 97.8% | 97.8% | 97.8% | 97.8% |
| **Duplicate** | 62.5% | 1.1% | 1.3% | 1.3% | 1.3% |
| **Fragment** | 0.7% | 0.7% | 0.6% | 0.6% | 0.6% |
| **Missing** | 0.4% | 0.4% | 0.3% | 0.3% | 0.3% |
| **n** | 2,510 | 2,510 | 2,510 | 2,510 | 2,510 |

### Quality, completeness, and coverage

Keeping in mind that accuracy of an assembly without an existing reference for a species is difficult to assess, our assembly appears to be high quality. At 1.162 Gbp with no gaps, the *T. sanguisuga* genome falls within an expected size range relative to *R. prolixus* (706.8 Mbp) and sister species *T. infestans* (949 Mbp; Table 3), suggesting a high level of completeness. The *R. prolixus* genome was the first triatomine species genome ever assembled, in 2015. Given advances in the field during the past ten years, the *R. prolixus* genome published had over 142 million unknown bases (Ns), while our genome did not have any unknown bases, which might explain some of the size difference. The GC percentage was similar between species, varying by less than half a percentage point. Coverage was higher for *T. sanguisuga* than *R. prolixus* (77.7x vs 8.3x, respectively, Table 2) but lower than *T. infestans* (200.0x). Our assembly is the most contiguous of the three species (183 contigs for *T. sanguisuga* vs. 14,951 and 16,511 for *T. infestans* and *R. prolixus*, respectively), with an N50 that is roughly 87 times larger than *R. prolixus* and 873 times larger than *T. infestans* (Table 2). At 99.1%, our BUSCO score indicates excellent resolution compared to that of the other two *Triatominae* genomes (Figure 3).

**Table 3.** Comparison of quantitative genome characteristics between *T. sanguisuga* (from this study), *T. infestans* and *R. prolixus*.

|  | *T. sanguisuga* | *T. infestans* | *R. prolixus* |
| --- | --- | --- | --- |
| **Summary statistics** |  |  |  |
| Contigs | 183 | 14,951 | 16,511 |
| Contigs >5000bp | 183 | 14,849 | 4,598 |
| Contigs >10,000bp | 182 | 13,959 | 3,097 |
| Contigs >25,000bp | 151 | 9,815 | 1,862 |
| Contigs >50,000bp | 48 | 5,919 | 1,089 |
| Largest contig | 144,379,499 bp | 1,054,224 bp | 13,425,595 |
| Total Length | 1,162 Mb | 949Mb | 707Mb |
| N50 | 94,972,618 | 108,830 | 1,088,772 |
| GCs % | 33.56 | 34.03 | 33.94 |
| Coverage | 77.7x | 200.0x | 8.3x |
| **Mismatches** |  |  |  |
| #Ns per 100kbp | 0 | 342.49 | 20,118 |
| #Ns | 0 | 3,251,881 | 142,194,158 |

**Figure 3.**
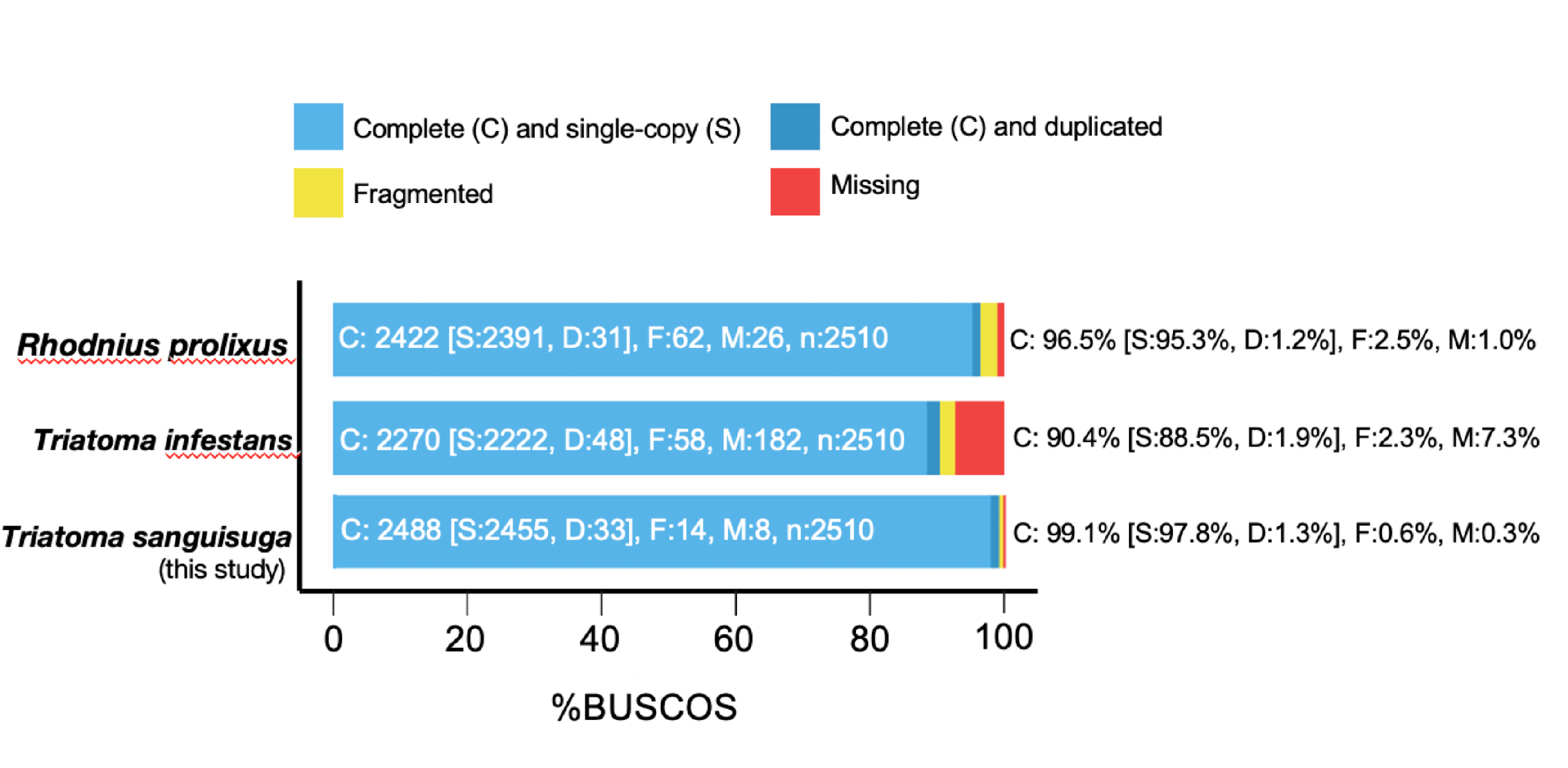
Genome BUSCO scores for assembled *Triatominae* genomes on NCBI. *Triatoma sanguisuga* shown is from this study.

### Genome annotation

We detected 17,799 putative protein-coding genes using the Maker genome annotation pipeline (Table 4). The final protein BUSCO score was 94.4% complete (92.4% single copy and 2.0% duplicates). This number is comparable to that of *R. prolixus*, which was predicted to have 15,456 protein-coding genes. In an analysis of the *R. prolixus* reference proteins using the web service OrthoVenn3 (Sun *et al*. 2023), 9,652 overlapping orthologous protein clusters were found, with 413 non-overlapping clusters found in *T. sanguisuga* and 300 in *R. prolixus* (Fig. 4). Annotations for *T. infestans* were not available on NCBI or Vectorbase, so a comparison with this species was not possible.

**Table 4.**
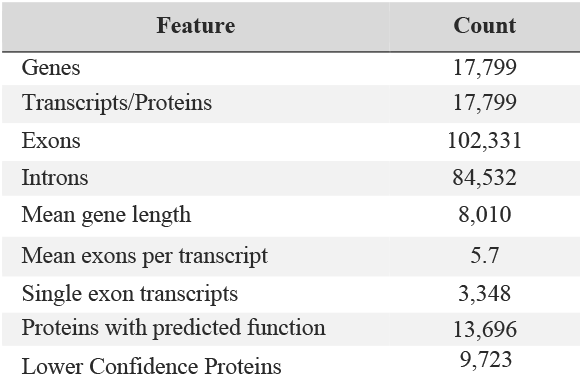
Quantitative summary of the annotated final genome of *T. sanguisuga*.

**Figure 4.**
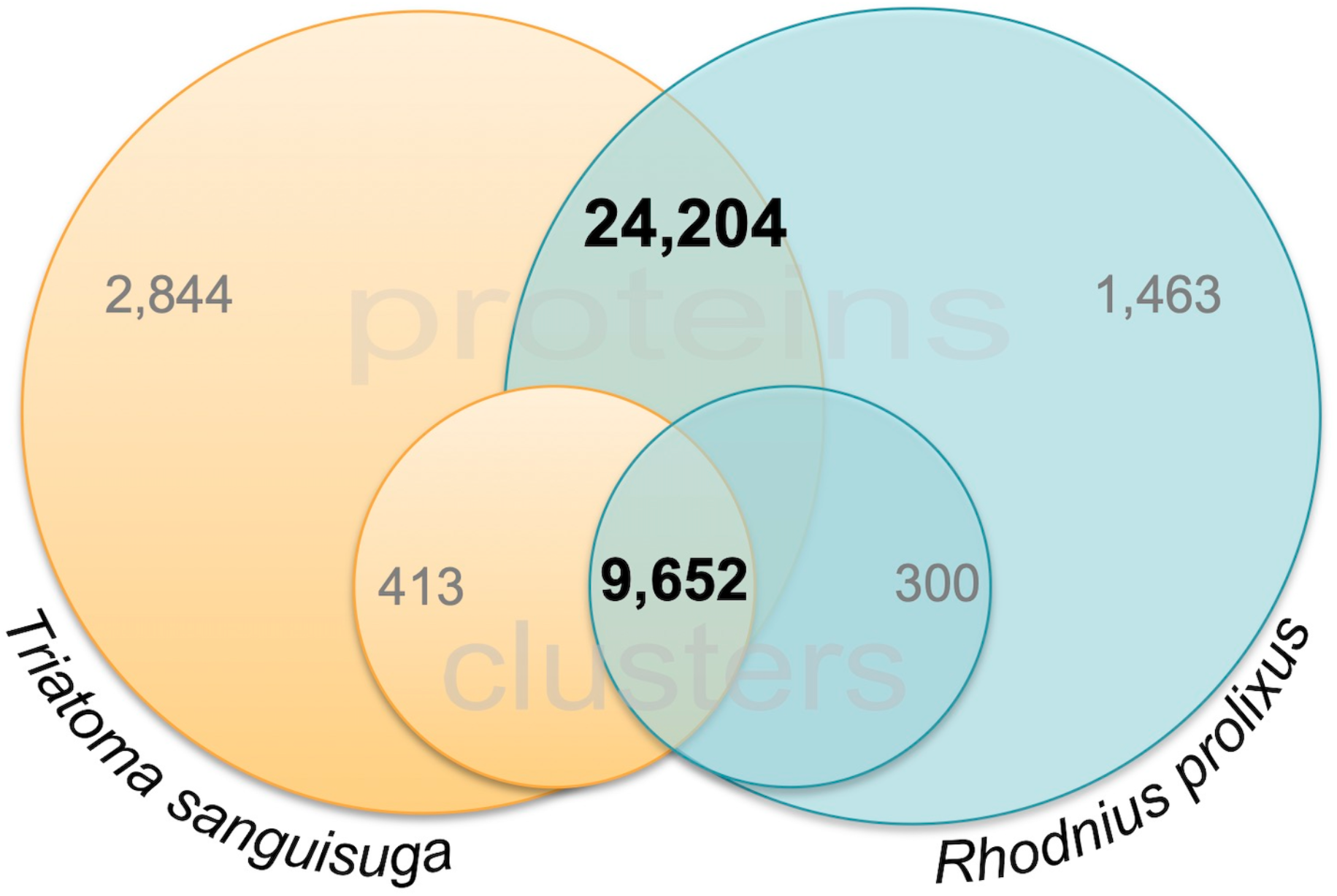
Protein count and cluster count overlap for *Triatoma sanguisuga* (orange) and *Rhodnius prolixus* (blue). Large circles indicate protein counts and small circles indicate orthologous protein cluster counts.

### Repetitive DNA

Repeat analysis using the RepeatModeler and RepeatMasker software revealed that the *T. sanguisuga* genome consists of 61.7% repetitive DNA, with 40.58% interspersed repeats, 2.35% simple repeats, and 0.46% low complexity repeats (Table 5).

**Table 5.** Interspersed repeats in the *T. sanguisuga* genome. *Most repeats fragmented by insertions or deletions have been counted as one element.

| Name | Number of elements* | Length | Percent |
| --- | --- | --- | --- |
| Retroelements | 869,571 | 214,680,453 bp | 18.47% |
| SINEs: | 66,050 | 6,690,116 bp | 0.58% |
| Penelope class | 88,144 | 115,173,646 bp | 1.31 % |
| LINE class: | 774,151 | 193,638,471 bp | 16.66% |
| CRE/SLACS | 0 | 0bp | 0.00% |
| L2/CR1/Rex | 34,854 | 20,442,282 bp | 1.76% |
| R1/LOA/Jockey | 390,559 | 77,649,217 bp | 6.68% |
| R2/R4/NeSL | 8,077 | 3,140,410 bp | 0.27% |
| RTE-Bov-B | 100,694 | 35,959,107 bp | 3.09% |
| L1/CIN4 | 91 | 49,474 bp | 0.00% |
| LTR elements: | 29,370 | 14,351,866 bp | 1.23% |
| BEL/Pao | 6,426 | 2,789,295 bp | 0.24% |
| Ty1/Copia | 8,706 | 1,639,481 bp | 0.14% |
| Gypsy/DIRS1 | 14,145 | 9,880,325 bp | 0.85% |
| Retroviral | 0 | 0 bp | 0.00% |
| DNA transposons | 509,635 | 84,165,895 bp | 7.24% |
| hobo-Activator | 50,682 | 15,582,243 bp | 1.34% |
| Tc1-IS630-Pogo | 154,823 | 27,948,296 bp | 2.40% |
| En-Spm | 0 | 0 bp | 0.00% |
| MuDR-IS905 | 0 | 0 bp | 0.00% |
| PiggyBac | 3,233 | 1,198,104 bp | 0.10% |
| Tourist/Harbinger | 2,099 | 674,150 bp | 0.06% |
| Other (Mirage, P-element, Transib) | 189 | 86,932 bp | 0.01% |
| Rolling-circles | 1,198,797 | 209,049,142 bp | 17.99% |
| Unclassified: | 649,059 | 169,616,531 bp | 14.60% |
| Total interspersed repeats: |  | 468,462,879 bp | 40.31% |
| Small RNA: | 24,593 | 4,690,086 bp | 0.40% |
| Satellites: | 7,833 | 1,134,749 bp | 0.10% |
| Simple repeats: | 693,752 | 27,246,855 bp | 2.34% |
| Low complexity: | 107,640 | 5,319,819 bp | 0.46% |

### Mitochondrial structure

Ninety-eight contigs from the original Hifiasm assembly were identified as redundant copies of the *T. sanguisuga* mitogenome. The representative mitochondrial contig selected and circularized by MitoHifi was aligned to the existing *T. sansguisuga* mitochondrial genome (NC_050329.1). BLAST hits between the representative contig and the existing *T. sanguisuga* mitochondrial sequence showed that there was an additional ∼1,500 bp in the representative contig. This approximate 1,500 bp was extracted and queried against nt and Univec build 10 databases. With no significant hits to either database, the region was subsequently removed from the contig. The final annotated mitogenome assembly was a single circular contig measuring 15,542 bp in length with 14 protein-coding genes, 2 rRNAs, and 22 tRNAs (Fig. 5). The sequence matched the reference sequence with 95.49% identity over 99% of its length.

**Figure 5.**
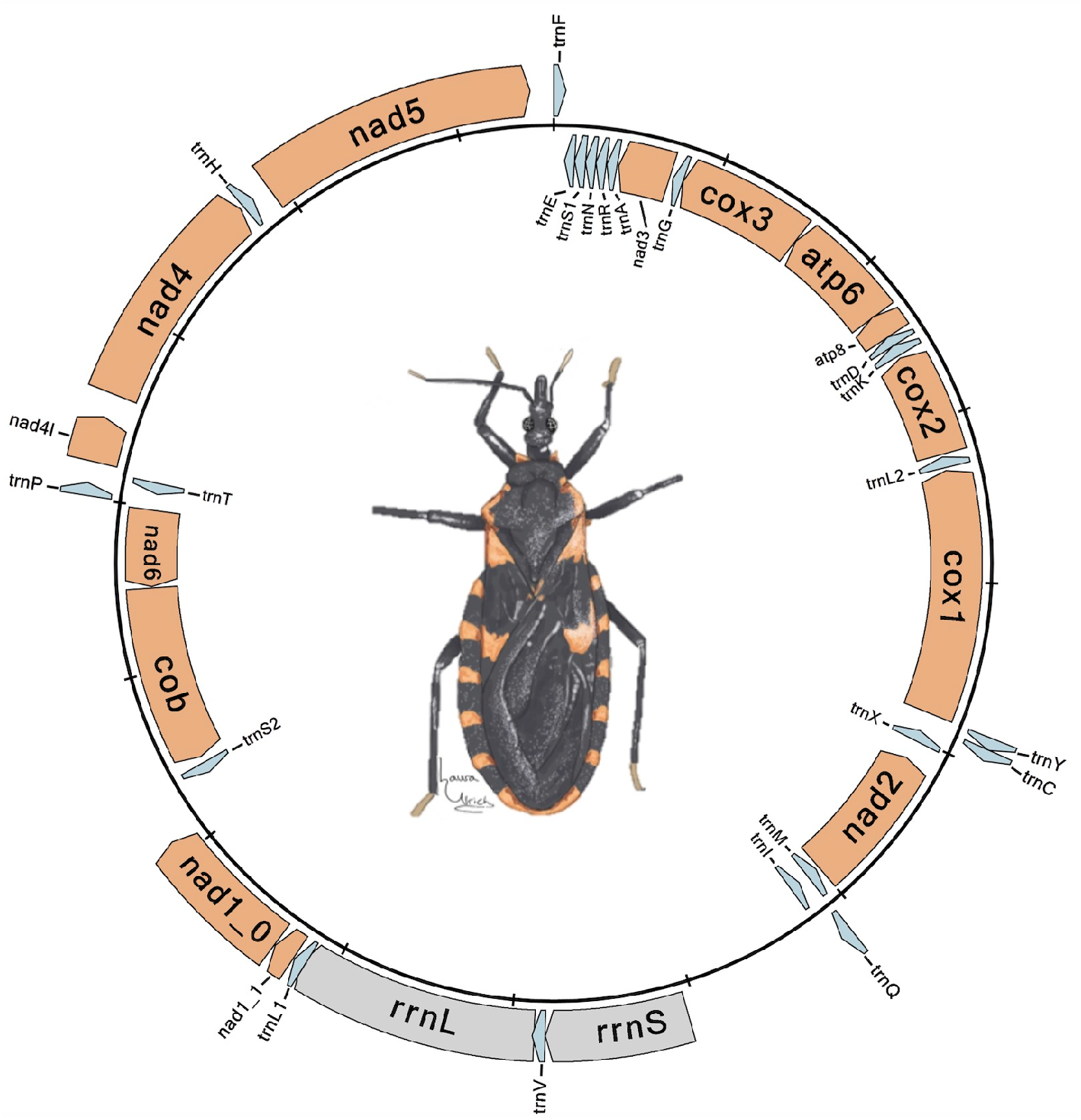
Fig. 5. Annotated mitogenome for *Triatoma sanguisuga*. Protein coding genes are shown in orange, tRNAs are shown in light blue, and rRNAs are shown in gray. Mitogenome was drawn using circularMT (Goodman and Carr 2024). Drawing of *T. sanguisuga* by Laura Ulrich (used with Ms. Ulrich’s permission.)

## IV. Conclusions

Here, we present the first genome sequence for *Triatoma sanguisuga*, which is also the first whole genome sequence for any US or North American triatomine species. The *T. sanguisuga* genome will facilitate the identification of genetic differences between *T. sanguisuga* populations, plausibly with relevance to epidemiologically important traits. Vector reference genomes contribute to our ability to carry out genetic investigations of physiological and behavioral attributes of disease vectors such as blood feeding, host seeking, and parasite competence. Findings from such studies can be used to guide vector-borne disease management strategies and in turn, strengthen public health preparedness.

## IV. Data Availability Statement

Supplementary file one (S1), submitted with this manuscript, contains the full commands for all bioinformatics steps. Supplementary file 2 (S2) is available on GSA FigShare and contains the annotation files. Sequence data and the genome assembly are publicly available on NCBI under BioProject ID PRJNA1140168 and Accession number SRR2998870.

## V. Acknowledgments

We are grateful to Brewster Kingham, Erin Bernberg, and Olga Shevchenko from the University of Delaware Sequencing and Genotyping Center for assistance with experimental design, sample processing methods, and PacBio sequencing. We would also like to thank Shawn Polson from the University of Delaware Bioinformatics Data Science Core Facility and the Delaware homeowner who provided the *Triatoma sanguisuga* sample.

## VI. Conflict of Interest

None

## VII. Funder Information

Access to the University of Delaware Sequencing and Bioinformatics Core Centers was supported by the Institutional Development Award (IDeA) from the National Institute of Health’s National Institute of General Medical Sciences, under grant number P20GM103446. Support from the University of Delaware CBCB Bioinformatics Data Science Core Facility (RRID:SCR_017696) including use of the BIOMIX and BioStore computational resources was made possible through funding from Delaware INBRE (NIGMS P20 GM103446), NIH Shared Instrumentation Grant (S10 OD028725), the State of Delaware, and the Delaware Biotechnology Institute.

## Notes

### Competing Interest Statement

The authors have declared no competing interest.

