## Supplementary file 1 for "Genome report: First whole genome sequence of *Triatoma sanguisuga* (Le Conte, 1855), vector of Chagas disease"

### T. Sanguisuga Genome Assembly and Annotation

#### Raw Read QC

Quality of the raw PacBio HiFi reads was examined using NanoPlot v1.43.0.

```
READS="[path_to_hifi_reads]/m84193_240403_193205_s2.fastq"
PREFIX="Triatoma_sanguisuga"
OUTDIR="[path_to_hifi_reads]/raw_read_qc"

NanoPlot --threads 32  --outdir $OUTDIR --prefix $PREFIX --fastq $READS
```

#### Assembly

The HiFi reads were assembled by several different assembly tools: hifiasm, flye, and smrtlink.

##### Hifiasm

Hifiasm v0.19.5 assembly scripts

```
OUT="[path_to_assembly_directory]/hifiasm/triatoma_sanguisuga"
INFILES="[path_to_hifi_reads]/m84193_240403_193205_s2.fastq"

mkdir -p ${OUT}
hifiasm -o ${OUT} -t ${SLURM_CPUS_PER_TASK} ${INFILES}
hifiasm -o ${OUT} -t ${SLURM_CPUS_PER_TASK --write-paf --write-ec /dev/null
```

##### Flye

Flye v2.9 assembly scripts

```
OUTDIR=[path_to_assembly_directory]/flye
INFILES="[path_to_hifi_reads]/m84193_240403_193205_s2.fastq"
GENOME_SIZE=1g

flye --pacbio-hifi ${INFILES} --out-dir ${OUTDIR} --threads ${SLURM_CPUS_PER_TASK} --
genome-size ${GENOME_SIZE} --iterations 3
```

##### SMRTLINK

The phased and unphased SMRTLink assemblies were run on the SMRT Link portal v13.1.0. Default settings were used for the phased assembly while the unphased assembly had the phasing turned off.

#### Assembly Quality Assessment

QUAST v5.1.0 and BUSCO v5.4.7 were used to assess the completeness of each assembly and compare them to the genome assemblies of two closely related species, Rhodnius prolixus (GCA_000181055.3) and Triatoma infestans (GCA_011037195.1)

QUAST scripts

```
BASEDIR=[path_to_assembly_directory]
SMRTLINK_UNPHASED="${BASEDIR}/smrtlink/unphased/final_purged_primary_unphased.fasta"
SMRTLINK_PHASED="${BASEDIR}/smrtlink/phased/final_purged_primary_phased.fasta"
FLYE="${BASEDIR}/flye/assembly.fasta"
HIFIASM="${BASEDIR}/hifiasm/triatoma_sanguisuga_hifiasm.bp.p_ctg.fa"
RHODNIUS_PROLIXUS="[path_to_reference_directory]/Rhodnius_prolixus_ref/VectorBase-
68_RprolixusCDC_Genome.fasta"
TRIATOMA_INFESTANS="
[path_to_reference_directory]/Triatoma_infestans_ref/GCA_011037195.1_UVM_Tinf_1.0_geno
mic.fna"

quast.py ${SMRTLINK_UNPHASED} ${SMRTLINK_PHASED} ${FLYE} ${HIFIASM}
${RHODNIUS_PROLIXUS} ${TRIATOMA_INFESTANS} \
  --min-contig 1000 -e --large -t 64 --labels "SMRTLINK UNPHASED","SMRTLINK
PHASED","FLYE","HIFIASM","RHODNIUS PROLIXUS","TRIATOMA INFESTANS" \
    -o quast_results
```

BUSCO scripts

```
export PATH=$PATH:/usr/local/bbmap
export NUMEXPR_MAX_THREADS=32

BASEDIR=[path_to_assembly_directory]
SMRTLINK_UNPHASED="${BASEDIR}/smrtlink/unphased/final_purged_primary_unphased.fasta"
SMRTLINK_PHASED="${BASEDIR}/smrtlink/phased/final_purged_primary_phased.fasta"
FLYE="${BASEDIR}/flye/assembly.fasta"
HIFIASM="${BASEDIR}/hifiasm/triatoma_sanguisuga_hifiasm.bp.p_ctg.fa"
HIFIASM_HAP1="${BASEDIR}/hifiasm/triatoma_sanguisuga_hifiasm.bp.hap1.p_ctg.fa"
RHODNIUS_PROLIXUS="[path_to_reference_directory]/Rhodnius_prolixus_ref/VectorBase-
68_RprolixusCDC_Genome.fasta"
TRIATOMA_INFESTANS="
[path_to_reference_directory]/Triatoma_infestans_ref/GCA_011037195.1_UVM_Tinf_1.0_geno
mic.fna"

SAMPLE="triatoma_sanguisuga"
LINEAGE="hemiptera_odb10"

busco -m genome -i ${FLYE} -o ${SAMPLE}_flye_busco -l ${LINEAGE} -c 32 -f
```

```
busco -m genome -i ${HIFIASM} -o ${SAMPLE}_hifiasm_busco -l ${LINEAGE} -c 32 -f
busco -m genome -i ${SMRTLINK_PHASED} -o ${SAMPLE}_smrtlink_phased_busco -l ${LINEAGE}
-c 32 -f
busco -m genome -i ${SMRTLINK_UNPHASED} -o ${SAMPLE}_smrtlink_unphased_busco -l
${LINEAGE} -c 32 -f
busco -m genome -i ${RHODNIUS_PROLIXUS} -o rhodnius_prolixus_busco -l ${LINEAGE} -c 32
-f
busco -m genome -i ${TRIATOMA_INFESTANS} -o triatoma_infestans_busco -l ${LINEAGE} -c
32 -f
```

#### Decontamination

BLASTn v2.15.0+ was used to query the contigs against the nt_core database v1.1 to check for contamination.

```
blastn -max_target_seqs 4 -evalue 0.01 -db core_nt -query
triatoma_sanguisuga_hifiasm.bp.p_ctg.fa \
  -num_threads 24 -outfmt "6 qseqid sseqid pident length mismatch gapopen qstart qend
sstart send evalue bitscore qcovs staxids" \
  > blast_hifiasm_to_core_nt_v6_25_24.txt

awk '$4>=200' blast_hifiasm_to_core_nt_v6_25_24.txt >
blast_hifiasm_to_core_nt_v6_25_24.filt200bp.txt
```

The taxon IDs in the last column of the blast output were used to check for contamination. Contigs with only hits to insects and ribosomal RNA were not considered contamination. QUAST was rerun (same parameters as above) with the decontaminated assembly included.

#### Mitochondria Identification

First, BLAST was used to query the contigs against the existing mitochondrial sequence of *T. sansuisuga* (NC_050329.1) downloaded from NCBI. Hits were filtered by query coverage to keep contigs with more >50% aligning to the mitochondrial sequences. This identified 98 contigs as possible mitochondria sequence.

```
blastn -evalue 0.001 -query
[path_to_assembly_directory]/hifiasm/triatoma_sanguisuga_assembly.decontam.fa \
  -db NC_050329.1.fasta -num_threads 24 -outfmt "6 qseqid sseqid pident length
mismatch gapopen qstart qend sstart send evalue bitscore qcovs staxids" \
  > blast_hifiasm_to_existing_mito.txt

awk '$13>=50' blast_hifiasm_to_existing_mito.txt > poss_mito_contigs_w_ref.txt
```

We also ran MitoHifi v3.2.2 via its Singularity container.

```
export APPTAINER_BIND="[work_directory]/circularize:/[work_directory]/circularize"

singularity exec docker://ghcr.io/marcelauliano/mitohifi:master ./mitohifi.sh
```

The mitohifi.sh script contained:

```
export JAVA_HOME=/usr/lib/jvm/java-11-openjdk-amd64/
export PATH=$JAVA_HOME/bin:$PATH

findMitoReference.py --species "Triatoma sanguisuga" --type mitochondrion --outfolder
related_mito

mitohifi.py -c ../triatoma_sanguisuga_hifiasm.bp.p_ctg.fa -f
../related_mito/NC_050329.1.fasta -g ../related_mito/NC_050329.1.gb -t 32 --mitos
```

MitoHifi identifed the same 98 contigs identified by the BLAST search as possible circular mitochondria sequences. MitoHifi selected the representative mitochondria contig and modified it to start at tRNA-Phe. In the BLAST results described above, the representative contig had an approximately ~1500 base pair stretch of sequence that did not align to the existing mitochondrial sequence. This ~1500 bp sequence was extracted based on the blast coordinates and queried against the nt and UniVec databses.

```
bedtools getfasta -fi MitoHifi_output/final_mitogenome_renamed.fasta -bed gap.bed -fo
gap.fa

#used NCBI website BLAST to query 'gap.fa' against NT (July 2024)
#downloaded UniVec database build 10 from https://ftp.ncbi.nlm.nih.gov/pub/UniVec/

makeblastdb -in UniVec.fa -dbtype nucl

blastn -query gap.fa -db UniVec.fa -outfmt 6 > gap_vs_univec.tsv

#the coordinates in 'nogap.bed' were identifed from the BLAST alignment to the
existing T sanguisuga mitochondrial sequence

bedtools getfasta -fi MitoHifi_output/final_mitogenome.fasta -bed nongap.bed -fo
mitochondria_contig99_nogap.fa
```

QUAST was rerun (same parameters as in the 'Assembly Quality Assessment' section) on the assembly with the redundant mitochondrial contigs removed and with the updated representative mitochondria contig included. MitoHifi was rerun on mitochondria_contig99_nogap.fa for annotation and the resulting GFF file was given to CircularMT for visualization.

#### Methylation

The reads with 5mC information were downloaded from PacBio SMRTLink portal and aligned to the assembly using pbmm2. PacBio's CpG-tools v2.3.2 and 'awk' were used to calculated global methylation.

```
READS="./methylation/with_5mC.fq"

ASSEMBLY="
[path_to_assembly_directory]/hifiasm/triatoma_sanguisuga_hifiasm.bp.p_ctg.fa"
PREFIX="pbmm2_pacbio_reads_w_5mC_to_HIFIASM"

pbmm2 align triatoma_sanguisuga_hifiasm.bp.p_ctg.mmi with_5mC.bam
pbmm2_pacbio_reads_w_5mC_to_HIFIASM.bam

samtools view -@ 32 -b -S $PREFIX.sam > $PREFIX.bam
samtools sort -@ 32 $PREFIX.bam > $PREFIX.sort.bam
samtools stats -@ 32  $PREFIX.sort.bam > $PREFIX.sort.stats
amtools flagstat -@ 32 $PREFIX.sort.bam > $PREFIX.sort.flagstats
samtools index ${PREFIX}.sort.bam

~/bin/pb-CpG-tools-v2.3.2-x86_64-unknown-linux-gnu/bin/aligned_bam_to_cpg_scores --
pileup-mode count --bam pbmm2_pacbio_reads_w_5mC_to_HIFIASM.sort.bam

awk '$5>0 {nonmod+=$8; mod+=$7} END{print 100*(mod/(nonmod+mod))}'
aligned_bam_to_cpg_scores.combined.bed
```

### RepeatMasking

Repeat modeling and masking was done on the original hifiasm primary assembly as well the decontaminated draft of the assembly and then the final assembly with the redundant mitochondria contigs removed. Scripts below are for the original hifiasm assembly but commands were kept the same for the other two versions of the assembly.

RepeatModeler script

```
FASTA_IN="
[path_to_assembly_directory]/hifiasm/triatoma_sanguisuga_hifiasm.bp.p_ctg.fa"
PREFIX="kissing_bug"

srun --job-name="BuildDatabase" BuildDatabase -name ${PREFIX} -engine ncbi ${FASTA_IN}
srun --job-name="RepeatModeler" RepeatModeler -pa 96 -engine ncbi -database ${PREFIX}
2>&1 | tee repeatmodeler.log
```

RepeatMasker script

```
FASTA_IN="
[path_to_assembly_directory]/hifiasm/triatoma_sanguisuga_hifiasm.bp.p_ctg.fa"
FASTA_EXT=fa
```

```
ENGINE="rmblast"

## TOP LEVEL MASK SHOULD BE REPBASE
#Repbase Mask
MASK=Triatominae
MASK_FILE=Triatominae
MASK_TYPE="-species"
BASE=`basename ${FASTA_IN} .${FASTA_EXT}`
DIR="${MASK}_mask"

mkdir -p ${DIR}
srun --job-name="RepeatMasker-1" RepeatMasker -pa ${SLURM_CPUS_PER_TASK} -e ${ENGINE}
${MASK_TYPE} ${MASK_FILE} -x -gff -dir ${DIR} ${FASTA_IN}
rename ${BASE}.${FASTA_EXT}. ${BASE}.${MASK}. ${DIR}/*
FASTA_EXT="masked"
FASTA_IN=${DIR}/${BASE}.${MASK}.${FASTA_EXT}
CAT_FILE+=("${DIR}/${BASE}.${MASK}.cat.gz")

#Custom de novo mask
MASK=KissingBugCustom
MASK_FILE=../1-repeatmodeler/kissing_bug-families.fa
MASK_TYPE="-lib"
BASE=`basename ${FASTA_IN} .${FASTA_EXT}`
DIR="${MASK}_mask"
mkdir -p ${DIR}
srun --job-name="RepeatMasker-2" RepeatMasker -pa ${SLURM_CPUS_PER_TASK} -e ${ENGINE}
${MASK_TYPE} ${MASK_FILE} -x -gff -dir ${DIR} ${FASTA_IN}
rename ${BASE}.${FASTA_EXT}. ${BASE}.${MASK}. ${DIR}/*
FASTA_IN=${DIR}/${BASE}.${MASK}.${FASTA_EXT}
OUT_FILE=${DIR}/${BASE}.${MASK}.out
CAT_FILE+=("${DIR}/${BASE}.${MASK}.cat.gz")

#Final
MASK=full
MASK_TYPE="-species"
BASE=`basename ${FASTA_IN} .${FASTA_EXT}`
DIR="${MASK}_mask"
mkdir -p ${DIR}
cp ${FASTA_IN} ${DIR}/${BASE}.${MASK}.${FASTA_EXT}
cp ${OUT_FILE} ${DIR}/${BASE}.${MASK}.orig.out
zcat `echo ${CAT_FILE[@]}` > ${DIR}/${BASE}.${MASK}.cat
srun --job-name="ProcessRepeats" ProcessRepeats ${MASK_TYPE} ${MASK_FILE} -gff -x
${DIR}/${BASE}.${MASK}.cat

# create GFF3
srun --job-name="rmOutToGFF3" rmOutToGFF3.pl ${DIR}/${BASE}.${MASK}.out  >
${DIR}/${BASE}.${MASK}.gff3

# to feed these repeats into MAKER, we need to extract the complex repeats
grep -v -e "Satellite" -e ")n" -e "-rich" ${DIR}/${BASE}.${MASK}.gff3 >
${DIR}/${BASE}.${MASK}.complex.gff3

# reformat to work with MAKER
cat ${DIR}/${BASE}.${MASK}.complex.gff3 | \
  perl -ane '$id; if(!/^\#/){@F = split(/\t/, $_); chomp $F[-1];$id++; $F[-1] .=
```

```
"\;ID=$id"; $_ = join("\t", @F)."\n"} print $_' \
  > ${DIR}/${BASE}.${MASK}.complex.reformat.gff3

ln -s ${DIR}/${BASE}.${MASK}.complex.reformat.gff3 ${BASE}.final.gff3
```

The commands for converting RepeatMasker's results to a format that works with Maker is based on code from the 'Repeat Annotation' section part 2 of 'Genome Annotation using MAKER' (https://gist.github.com/darencard/bb1001ac1532dd4225b030cf0cd61ce2).

### Annotation

#### Maker Round 1

The first, evidence based round of Maker was run using the transcripts from Rhodnius prolixus and the 'Triatominae' canonical proteins from UniProt release 2024_3 as the transcript and protein evidence respectively. For the repeat masking, the 'rm_gff' parameter was set to the complex repeats GFF from RepeatMasker (see commands above).

```
source ../maker-pipe.source

RD=1
STUB=${BASE}-rd${RD}
SLOTS=$(( ${SLURM_CPUS_PER_TASK} * ${SLURM_NTASKS} ))

mpirun -n ${SLOTS} maker -fix_nucleotides -base ${STUB} maker_opts.ctl maker_bopts.ctl
maker_exe.ctl

cd ${STUB}.maker.output
srun --ntasks=1 --job-name="gff3_merge-full" gff3_merge -s -d
${STUB}_master_datastore_index.log > ${STUB}.all.maker.gff
srun --ntasks=1 --job-name="fasta_mege" fasta_merge -d
${STUB}_master_datastore_index.log
# GFF w/o the sequences
srun --ntasks=1 --job-name="gff3_merge-noseq" gff3_merge -n -s -d
${STUB}_master_datastore_index.log > ${STUB}.all.maker.noseq.gff
cd ..
```

The maker_opts.ctl file contained:

```
#-----Genome (these are always required)
genome=[path_to_assembly_directory]/hifiasm/triatoma_sanguisuga_hifiasm.bp.p_ctg.fa
organism_type=eukaryotic #eukaryotic or prokaryotic. Default is eukaryotic

#-----Re-annotation Using MAKER Derived GFF3
maker_gff= #MAKER derived GFF3 file
est_pass=0 #use ESTs in maker_gff: 1 = yes, 0 = no
altest_pass=0 #use alternate organism ESTs in maker_gff: 1 = yes, 0 = no
protein_pass=0 #use protein alignments in maker_gff: 1 = yes, 0 = no
rm_pass=0 #use repeats in maker_gff: 1 = yes, 0 = no
```

```
model_pass=0 #use gene models in maker_gff: 1 = yes, 0 = no
pred_pass=0 #use ab-initio predictions in maker_gff: 1 = yes, 0 = no
other_pass=0 #passthrough anyything else in maker_gff: 1 = yes, 0 = no

#-----EST Evidence (for best results provide a file for at least one)
est=[path_to_reference_directory]/Rhodnius_prolixus_ref/VectorBase-
68_RprolixusCDC_AnnotatedTranscripts.fasta
altest=
est_gff= #aligned ESTs or mRNA-seq from an external GFF3 file
altest_gff= #aligned ESTs from a closly relate species in GFF3 format

#-----Protein Homology Evidence (for best results provide a file for at least one)
protein=[path_to_annotation_directory]/uniprotkb_Triatominae_2024_05_28.fasta
protein_gff=

#-----Repeat Masking (leave values blank to skip repeat masking)
model_org=simple #select a model organism for RepBase masking in RepeatMasker
rmlib=
repeat_protein=te_proteins.fasta #provide a fasta file of transposable element
proteins for RepeatRunner
rm_gff=[path_to_annotation_directory]/2-
repeatmasker/triatoma_sanguisuga_hifiasm.bp.p_ctg.Triatominae.KissingBugCustom.final.g
ff3
prok_rm=0 #forces MAKER to repeatmask prokaryotes (no reason to change this), 1 = yes,
0 = no
softmask=1 #use soft-masking rather than hard-masking in BLAST (i.e. seg and dust
filtering)

#-----Gene Prediction
snaphmm= #SNAP HMM file
gmhmm= #GeneMark HMM file
augustus_species= #Augustus gene prediction species model
fgenesh_par_file= #FGENESH parameter file
pred_gff= #ab-initio predictions from an external GFF3 file
model_gff= #annotated gene models from an external GFF3 file (annotation pass-through)
run_evm=0 #run EvidenceModeler, 1 = yes, 0 = no
est2genome=1 #infer gene predictions directly from ESTs, 1 = yes, 0 = no
protein2genome=1 #infer predictions from protein homology, 1 = yes, 0 = no
trna=0 #find tRNAs with tRNAscan, 1 = yes, 0 = no
snoscan_rrna= #rRNA file to have Snoscan find snoRNAs
snoscan_meth= #-O-methylation site fileto have Snoscan find snoRNAs
unmask=0 #also run ab-initio prediction programs on unmasked sequence, 1 = yes, 0 = no
allow_overlap= #allowed gene overlap fraction (value from 0 to 1, blank for default)

#-----Other Annotation Feature Types (features MAKER doesn't recognize)
other_gff= #extra features to pass-through to final MAKER generated GFF3 file

#-----External Application Behavior Options
alt_peptide=C #amino acid used to replace non-standard amino acids in BLAST databases
cpus=1 #max number of cpus to use in BLAST and RepeatMasker (not for MPI, leave 1 when
using MPI)

#-----MAKER Behavior Options
max_dna_len=100000 #length for dividing up contigs into chunks (increases/decreases
memory usage)
```

```
min_contig=1 #skip genome contigs below this length (under 10kb are often useless)

pred_flank=200 #flank for extending evidence clusters sent to gene predictors
pred_stats=0 #report AED and QI statistics for all predictions as well as models
AED_threshold=1 #Maximum Annotation Edit Distance allowed (bound by 0 and 1)
min_protein=0 #require at least this many amino acids in predicted proteins
alt_splice=0 #Take extra steps to try and find alternative splicing, 1 = yes, 0 = no
always_complete=0 #extra steps to force start and stop codons, 1 = yes, 0 = no
map_forward=0 #map names and attributes forward from old GFF3 genes, 1 = yes, 0 = no
keep_preds=1 #Concordance threshold to add unsupported gene prediction (bound by 0 and
1)

split_hit=10000 #length for the splitting of hits (expected max intron size for
evidence alignments)
min_intron=20 #minimum intron length (used for alignment polishing)
single_exon=0 #consider single exon EST evidence when generating annotations, 1 = yes,
0 = no
single_length=250 #min length required for single exon ESTs if 'single_exon is
enabled'
correct_est_fusion=0 #limits use of ESTs in annotation to avoid fusion genes

tries=2 #number of times to try a contig if there is a failure for some reason
clean_try=0 #remove all data from previous run before retrying, 1 = yes, 0 = no
clean_up=0 #removes theVoid directory with individual analysis files, 1 = yes, 0 = no
TMP=/scratch/tmp #specify a directory other than the system default temporary
directory for temporary files
```

The maker_bopts.ctl file contained:

```
#-----BLAST and Exonerate Statistics Thresholds
blast_type=ncbi+ #set to 'ncbi+', 'ncbi' or 'wublast'
use_rapsearch=0 #use rapsearch instead of blastx, 1 = yes, 0 = no

pcov_blastn=0.8 #Blastn Percent Coverage Threhold EST-Genome Alignments
pid_blastn=0.85 #Blastn Percent Identity Threshold EST-Genome Aligments
eval_blastn=1e-10 #Blastn eval cutoff
bit_blastn=40 #Blastn bit cutoff
depth_blastn=20 #Blastn depth cutoff (0 to disable cutoff)

pcov_blastx=0.5 #Blastx Percent Coverage Threhold Protein-Genome Alignments
pid_blastx=0.4 #Blastx Percent Identity Threshold Protein-Genome Aligments
eval_blastx=1e-06 #Blastx eval cutoff
bit_blastx=30 #Blastx bit cutoff
depth_blastx=20 #Blastx depth cutoff (0 to disable cutoff)

pcov_tblastx=0.8 #tBlastx Percent Coverage Threhold alt-EST-Genome Alignments
pid_tblastx=0.85 #tBlastx Percent Identity Threshold alt-EST-Genome Aligments
eval_tblastx=1e-10 #tBlastx eval cutoff
bit_tblastx=40 #tBlastx bit cutoff
depth_tblastx=20 #tBlastx depth cutoff (0 to disable cutoff)

pcov_rm_blastx=0.5 #Blastx Percent Coverage Threhold For Transposable Element Masking
pid_rm_blastx=0.4 #Blastx Percent Identity Threshold For Transposbale Element Masking
```

```
eval_rm_blastx=1e-06 #Blastx eval cutoff for transposable element masking
bit_rm_blastx=30 #Blastx bit cutoff for transposable element masking

ep_score_limit=20 #Exonerate protein percent of maximal score threshold
en_score_limit=20 #Exonerate nucleotide percent of maximal score threshold
```

#### Augustus Training

BUSCO was used to train Augustus using the transcripts from the 1st round of maker and 'rhodnius' as the starting model species.

```
source ../maker-pipe.source

export NUMEXPR_MAX_THREADS=8

RD=1
MAKER_DIR=${BASE_DIR}/3-maker-rd1
STUB=${BASE}-rd${RD}
MAKER_NOSEQ_GFF="[path_to_annotation_directory]/3-maker-rd1/Triatoma_sanguisuga-
rd1.maker.output/Triatoma_sanguisuga-rd1.all.maker.noseq.gff"

srun --job-name="generate_transcripts" awk -v OFS="\t" '{ if ($3 == "mRNA") print $1,
$4, $5 }' ${MAKER_NOSEQ_GFF} | \
awk -v OFS="\t" '{ if ($2 < 1000) print $1, "0", $3+1000; else print $1, $2-1000,
$3+1000 }' | \
bedtools getfasta -fi ${FASTA_IN} -bed - -fo ${STUB}.all.maker.transcripts1000.fasta

busco -l ${BUSCO_LIN} --restart -i ${STUB}.all.maker.transcripts1000.fasta -o ${STUB}
-m genome -c 8 --long --augustus --augustus_species="rhodnius" --
augustus_parameters='--progress=true'

# RENAME AND COPY TO AUGUSTUS CONFIG
cd ${STUB}/run*/augustus_output/retraining_parameters/BUSCO_${STUB}
rename BUSCO_${STUB} ${STUB} *
sed -i "s/BUSCO_${STUB}/${STUB}/g" ${STUB}_parameters.cfg
sed -i "s/BUSCO_${BASE}/${STUB}/g" ${STUB}_parameters.cfg.orig1
mkdir -p $AUGUSTUS_CONFIG_PATH/species/${STUB}
cp ${STUB}* $AUGUSTUS_CONFIG_PATH/species/${STUB}/
```

#### Maker Round 2

The second round of maker used the trained augustus model from the above commands.

```
source ../maker-pipe.source

RD=2
PREV_DIR=../3-maker-rd1
STUB=${BASE}-rd${RD}
PREV_RD=$(($RD-1))
PREV_STUB=${BASE}-rd${PREV_RD}
```

```
MAKER_NOSEQ_GFF=${PREV_DIR}/${PREV_STUB}.maker.output/${PREV_STUB}.all.maker.noseq.gff
SLOTS=$(( ${SLURM_CPUS_PER_TASK} * ${SLURM_NTASKS} ))

# transcript alignments
awk '{ if ($2 == "est2genome") print $0 }' ${MAKER_NOSEQ_GFF} >
${PREV_DIR}/${PREV_STUB}.all.maker.est2genome.gff
# protein alignments
awk '{ if ($2 == "protein2genome") print $0 }' ${MAKER_NOSEQ_GFF} >
${PREV_DIR}/${PREV_STUB}.all.maker.protein2genome.gff
# repeat alignments
awk '{ if ($2 ~ "repeat") print $0 }' ${MAKER_NOSEQ_GFF} >
${PREV_DIR}/${PREV_STUB}.all.maker.repeats.gff
# cdna(altest) alignments
awk '{ if ($2 == "cdna2genome") print $0 }' ${MAKER_NOSEQ_GFF} >
${PREV_DIR}/${PREV_STUB}.all.maker.cdna2genome.gff

#MAKER Command
mpirun -n ${SLOTS} maker -fix_nucleotides -base ${STUB} maker_opts.ctl maker_bopts.ctl
maker_exe.ctl

cd ${STUB}.maker.output
srun --ntasks=1 --job-name="gff3_merge-full" gff3_merge -s -d
${STUB}_master_datastore_index.log > ${STUB}.all.maker.gff
srun --ntasks=1 --job-name="fasta_merge" fasta_merge -d
${STUB}_master_datastore_index.log
# GFF w/o the sequences
srun --ntasks=1 --job-name="gff3_merge-noseq" gff3_merge -n -s -d
${STUB}_master_datastore_index.log > ${STUB}.all.maker.noseq.gff

# AED SUMMARY
AED_cdf_generator.pl -b 0.025 ${STUB}.all.maker.gff > ${STUB}.aed.txt
cd ..

#BUSCO SUMMARY
srun --ntasks=1 --job-name="busco_tran" busco -f -i
${STUB}.maker.output/${STUB}.all.maker.transcripts.fasta -o BUSCO-tran-${STUB} -m
transcriptome -l ${BUSCO_LIN} -c ${SLOTS} --update-data
srun --ntasks=1 --job-name="generate_plot_tran" generate_plot.py -wd BUSCO-
tran-${STUB}
rename busco_figure ${STUB}-busco-transcript-summary BUSCO-tran-${STUB}/*

srun --ntasks=1 --job-name="busco_prot" busco -f -i
${STUB}.maker.output/${STUB}.all.maker.proteins.fasta -o BUSCO-prot-${STUB} -m
proteins -l ${BUSCO_LIN} -c ${SLOTS}
srun --ntasks=1 --job-name="generate_plot_prot" generate_plot.py -wd BUSCO-
prot-${STUB}
rename busco_figure ${STUB}-busco-protein-summary BUSCO-prot-${STUB}/*
```

The maker_opts.ctl file contained:

```
#-----Genome (these are always required)
genome=[path_to_assembly_directory]/triatoma_sanguisuga_hifiasm.bp.p_ctg.fa
organism_type=eukaryotic #eukaryotic or prokaryotic. Default is eukaryotic
```

```
#-----Re-annotation Using MAKER Derived GFF3
maker_gff= #MAKER derived GFF3 file
est_pass=0 #use ESTs in maker_gff: 1 = yes, 0 = no
altest_pass=0 #use alternate organism ESTs in maker_gff: 1 = yes, 0 = no
protein_pass=0 #use protein alignments in maker_gff: 1 = yes, 0 = no
rm_pass=0 #use repeats in maker_gff: 1 = yes, 0 = no
model_pass=0 #use gene models in maker_gff: 1 = yes, 0 = no
pred_pass=0 #use ab-initio predictions in maker_gff: 1 = yes, 0 = no
other_pass=0 #passthrough anyything else in maker_gff: 1 = yes, 0 = no

#-----EST Evidence (for best results provide a file for at least one)
est=
altest= #EST/cDNA sequence file in fasta format from an alternate organism
est_gff=[path_to_annoation_directory]/3-maker-rd1/Triatoma_sanguisuga-
rd1.all.maker.est2genome.gff #aligned ESTs or mRNA-seq from an external GFF3 file
altest_gff=

#-----Protein Homology Evidence (for best results provide a file for at least one)
protein=
protein_gff=[path_to_annoation_directory]/3-maker-rd1/Triatoma_sanguisuga-
rd1.all.maker.protein2genome.gff

#-----Repeat Masking (leave values blank to skip repeat masking)
model_org= #select a model organism for RepBase masking in RepeatMasker
rmlib=
repeat_protein=te_proteins.fasta #provide a fasta file of transposable element
proteins for RepeatRunner
rm_gff=[path_to_annoation_directory]/3-maker-rd1/Triatoma_sanguisuga-
rd1.all.maker.repeats.gff
prok_rm=0 #forces MAKER to repeatmask prokaryotes (no reason to change this), 1 = yes,
0 = no
softmask=1 #use soft-masking rather than hard-masking in BLAST (i.e. seg and dust
filtering)

#-----Gene Prediction
snaphmm=
gmhmm= #GeneMark HMM file
augustus_species=Triatoma_sanguisuga-rd1
fgenesh_par_file= #FGENESH parameter file
pred_gff= #ab-initio predictions from an external GFF3 file
model_gff=
run_evm=0 #run EvidenceModeler, 1 = yes, 0 = no
est2genome=0 #infer gene predictions directly from ESTs, 1 = yes, 0 = no
protein2genome=0 #infer predictions from protein homology, 1 = yes, 0 = no
trna=0 #find tRNAs with tRNAscan, 1 = yes, 0 = no
snoscan_rrna= #rRNA file to have Snoscan find snoRNAs
snoscan_meth= #-O-methylation site fileto have Snoscan find snoRNAs
unmask=1 #also run ab-initio prediction programs on unmasked sequence, 1 = yes, 0 = no
allow_overlap= #allowed gene overlap fraction (value from 0 to 1, blank for default)

#-----Other Annotation Feature Types (features MAKER doesn't recognize)
other_gff= #extra features to pass-through to final MAKER generated GFF3 file

#-----External Application Behavior Options
```

```
alt_peptide=C #amino acid used to replace non-standard amino acids in BLAST databases
cpus=1 #max number of cpus to use in BLAST and RepeatMasker (not for MPI, leave 1 when
using MPI)

#-----MAKER Behavior Options
max_dna_len=100000 #length for dividing up contigs into chunks (increases/decreases
memory usage)
min_contig=1 #skip genome contigs below this length (under 10kb are often useless)

pred_flank=200 #flank for extending evidence clusters sent to gene predictors
pred_stats=0 #report AED and QI statistics for all predictions as well as models
AED_threshold=1 #Maximum Annotation Edit Distance allowed (bound by 0 and 1)
min_protein=0 #require at least this many amino acids in predicted proteins
alt_splice=0 #Take extra steps to try and find alternative splicing, 1 = yes, 0 = no
always_complete=0 #extra steps to force start and stop codons, 1 = yes, 0 = no
map_forward=0 #map names and attributes forward from old GFF3 genes, 1 = yes, 0 = no
keep_preds=1 #Concordance threshold to add unsupported gene prediction (bound by 0 and
1)

split_hit=10000 #length for the splitting of hits (expected max intron size for
evidence alignments)
min_intron=20 #minimum intron length (used for alignment polishing)
single_exon=0 #consider single exon EST evidence when generating annotations, 1 = yes,
0 = no
single_length=250 #min length required for single exon ESTs if 'single_exon is
enabled'
correct_est_fusion=0 #limits use of ESTs in annotation to avoid fusion genes

tries=2 #number of times to try a contig if there is a failure for some reason
clean_try=0 #remove all data from previous run before retrying, 1 = yes, 0 = no
clean_up=0 #removes theVoid directory with individual analysis files, 1 = yes, 0 = no
TMP=/scratch/tmp #specify a directory other than the system default temporary
directory for temporary files
```

The maker_bopts.ctl was identical to the one used in the first round of Maker.

#### Functional Assignment

Putative gene function was assigned by following Maker Support Protocols 2 and 3 (https://www.yandell-lab.org/publications/pdf/maker_current_protocols.pdf -Campbell et al. 2014).

```
GFF=Triatoma_sanguisuga-rd2.all.maker.noseq.gff
TRANSCRIPTS=Triatoma_sanguisuga-rd2.all.maker.transcripts.fasta
PROTEINS=Triatoma_sanguisuga-rd2.all.maker.proteins.fasta

[maker_install_dir]/bin/maker_map_ids --prefix TSAN_ --justify 6 $GFF >
Triatoma_sanguisuga-rd2.all.map

[maker_install_dir]/bin/map_fasta_ids Triatoma_sanguisuga-rd2.all.map $TRANSCRIPTS
[maker_install_dir]/bin/map_fasta_ids Triatoma_sanguisuga-rd2.all.map $PROTEINS
[maker_install_dir]/bin/map_gff_ids  Triatoma_sanguisuga-rd2.all.map $GFF
```

```
makeblastdb -in uniprot_sprot.fasta -input_type fasta -dbtype prot

blastp -db uniprot_sprot.fasta -query $PROTEINS -out maker2uni.blastp -evalue 0.000001
-outfmt 6 -num_alignments 1 -num_threads 16 \
  -seg yes -soft_masking true -lcase_masking -max_hsps 1

[maker_install_dir]/bin/maker_functional_gff uniprot_sprot.fasta maker2uni.blastp $GFF
> Triatoma_sanguisuga-rd2.all.maker.function.gff
[maker_install_dir]/bin/maker_functional_fasta uniprot_sprot.fasta maker2uni.blastp
$PROTEINS > Triatoma_sanguisuga-rd2.all.maker.proteins.function.fasta
[maker_install_dir]/bin/maker_functional_fasta uniprot_sprot.fasta maker2uni.blastp
$TRANSCRIPTS > Triatoma_sanguisuga-rd2.all.maker.transcripts.function.fasta
```

In addition, Interproscan v5.53-87.0 was run on the annotated proteins using steps 4 and 5 in Basic Protocol 5 (https://www.yandell-lab.org/publications/pdf/maker_current_protocols.pdf - Campbell et al. 2014).

```
GFF="Triatoma_sanguisuga-rd2.all.maker.function.gff"
TRANSCRIPTS="Triatoma_sanguisuga-rd2.all.maker.transcripts.function.fasta"
PROTEINS="Triatoma_sanguisuga-rd2.all.maker.proteins.function.fasta"

interproscan.sh -appl PfamA -iprlookup -goterms -f tsv -i $PROTEINS

[maker_install_dir]/bin/ipr_update_gff $GFF Triatoma_sanguisuga-
rd2.all.maker.proteins.function.fasta.tsv > Triatoma_sanguisuga-
rd2.all.maker.function_ipr.gff
```

#### Function and AED Filtering

The predicted genes were sorted into a high confidence set by filtering out genes with no function assigned and an AED of 1 (no transcript or protein evidence from Maker)

```
grep '>' Triatoma_sanguisuga-rd2.all.maker.proteins.function.fa | sed s/'>'/''/g >
Triatoma_sanguisuga-rd2.all.maker.proteins.function.txt

grep -v 'AED:1.00' Triatoma_sanguisuga-rd2.all.maker.proteins.function.txt >
AEDlessthan1.txt

grep 'AED:1.00' Triatoma_sanguisuga-rd2.all.maker.proteins.function.txt | grep
'Similar' > AEDof1wfunction.txt

cat AEDlessthan1.txt AEDof1wfunction.txt > Triatoma_sanguisuga-
rd2.all.maker.proteins.function.highconf.txt

faSomeRecords Triatoma_sanguisuga-rd2.all.maker.proteins.function.fa
Triatoma_sanguisuga-rd2.all.maker.proteins.function.highconf.txt Triatoma_sanguisuga-
rd2.all.maker.proteins.function.highconf.fa
```

The same IDs removed from Triatoma_sanguisuga-rd2.all.maker.proteins.function.fa to create Triatoma_sanguisuga-rd2.all.maker.proteins.function.highconf.txt were filtered out of the transcript file (faSomeRecords) and the GFF file (agat_sp_filter_feature_from_kill_list.pl via AGAT singularity v0.8.1).

#### Annotation Quality

BUSCO was run in protein and transcript mode on the high confidence proteins/transcripts from the annotation.

```
export PATH=$PATH:/usr/local/bbmap
export NUMEXPR_MAX_THREADS=32

PROTEINS="Triatoma_sanguisuga-rd2.all.maker.proteins.function.highconf.fasta"
TRANSCRIPTS="Triatoma_sanguisuga-rd2.all.maker.transcripts.function.highconf.fasta"
SAMPLE="Triatoma_sanguisuga"
LINEAGE="hemiptera_odb10"

busco -m transcriptome -i ${TRANSCRIPTS} -o ${SAMPLE}_transcript_busco -l ${LINEAGE} -
c 32 -f
busco -m protein -i ${PROTEINS} -o ${SAMPLE}_protein_busco -l ${LINEAGE} -c 32 -f
```
